# Early life stress reduces voluntary exercise and its prevention of diet-induced obesity and metabolic dysfunction in mice

**DOI:** 10.1101/2020.01.24.918805

**Authors:** Olivia C. Eller-Smith, E. Matthew Morris, John P. Thyfault, Julie A. Christianson

**Author notes:** Correspondence to: Julie Christianson, PhD, Department of Anatomy and Cell Biology, University of Kansas Medical Center, 3901 Rainbow Boulevard, MS 3038, Kansas City, KS 66160.

## Abstract

The development of obesity-related metabolic syndrome (MetS) involves a complex interaction of genetic and environmental factors. One environmental factor found to be significantly associated with MetS is early life stress (ELS). We have previously reported on our mouse model of ELS, induced by neonatal maternal separation (NMS), that displays altered regulation of the hypothalamic-pituitary-adrenal (HPA) axis and increased sensitivity in the urogenital organs, which was attenuated by voluntary wheel running. Here, we are using our NMS model to determine if ELS-induced changes in the HPA axis also influence weight gain and MetS. Naïve (non-stressed) and NMS male mice were given free access to a running wheel and a low-fat control diet at 4-weeks of age. At 16-weeks of age, half of the mice were transitioned to a high fat/sucrose (HFS) diet to investigate if NMS influences the effectiveness of voluntary exercise to prevent diet-induced obesity and MetS. Overall, we observed a greater impact of voluntary exercise on prevention of HFS diet-induced outcomes in naïve mice, compared to NMS mice. Although body weight and fat mass were still significantly higher, exercise attenuated fasting insulin levels and mRNA levels of inflammatory markers in epididymal adipose tissue in HFS diet-fed naïve mice. Only moderate changes were observed in exercised NMS mice on a HFS diet, although this could partially be explained by reduced running distance within this group. Interestingly, sedentary NMS mice on a control diet displayed impaired glucose homeostasis and moderately increased pro-inflammatory mRNA levels in epididymal adipose, suggesting that early life stress alone impairs metabolic function and negatively impacts the therapeutic effect of voluntary exercise.

## 1. Introduction

Stress is described as a change in homeostasis caused by various physiological, psychological, or environmental factors [1]. Although acute stress is necessary for survival, long-term stress, particularly in childhood, can lead to the development of chronic health disorders, including obesity [2–5]. Although the mechanism underlying this association has not been fully established, dysregulation of the hypothalamic-pituitary-adrenal (HPA) axis has been implicated. The HPA axis is a neuroendocrine system that contributes to the regulation of the stress response [6]. During a stressful event, the hypothalamus releases corticotropin-releasing factor (CRF), which signals the pituitary gland to release adrenocorticotrophic hormone (ACTH). ACTH then signals the adrenal cortex to release glucocorticoids, which include cortisol in humans and corticosterone in rodents [7]. Upon release, glucocorticoids modulate many downstream metabolic effects including alterations in glucose and fat metabolism as well as control of the cardiovascular system and the immune response [8]. CRF can also work in the periphery by activating mast cells (MC), which release cytokines and proteases that contribute to the formation of a pro-inflammatory environment [9, 10].

The HPA axis is not fully developed when children are born and therefore stressful events at this stage can cause long-term permanent damage on the regulation and output of this system, contributing to poor health outcomes in adulthood [11]. Childhood neglect and maltreatment are two forms of early life stress that are associated with the development of obesity in adulthood in humans [3–5]. Additionally, HPA axis hyperactivity is seen in patients with visceral obesity [12, 13]. Research in non-human primates [14] and rodents [15] also show early life stress-induced alterations in metabolic functions.

In addition to stress, other environmental risk factors associated with the development of obesity include a sedentary lifestyle and consumption of a ‘western diet’ (high proportion of saturated fat and refined sugars) [16–21]. This association is particularly troubling because sedentary behavior is highly prevalent in the United States, where most individuals do not meet the recommended requirements for daily physical activity [22] and western diet consumption has grown due to its convenience and low cost [23]. Visceral or central obesity is a component of the metabolic syndrome (MetS) [24], which is diagnosed when an individual meets 3 of the following 5 criteria: central obesity, insulin resistance, hypertension, high triglycerides, and low high-density lipoprotein cholesterol [25]. There is a high incidence of MetS in the United States, where 33% of adults meet the diagnostic criteria [26]. The constellation of risk factors associated with MetS synergizes to increase for both type 2 diabetes and cardiovascular disease.

It is clear that obesity and the subsequent development of MetS are substantial public health problems. Therefore, studying risk factors for the development of these chronic conditions as well as the most effective way to treat them is extremely important. In the present study, we use a model of early life stress in mice, neonatal maternal separation (NMS), that displays evidence of altered HPA axis regulation [27, 28] to test the hypothesis that early life stress-exposure increases the susceptibility to the detrimental metabolic consequences of the combination of a sedentary lifestyle and long-term western diet. Additionally, because lifestyle changes such as exercise and diet have been shown to improve symptoms of MetS in both humans [21, 29, 30] and rodents [31], we studied whether exercise, in the form of voluntary wheel running, could be used as a therapeutic intervention in the prevention of obesity-related MetS. To our knowledge, this is the first study to combine early life stress, diet, and exercise to study the development of factors related to MetS in mice.

## 2. Methods

### 2.1 Animals

All experiments in this study were performed on male C57Bl/6 mice (Charles River, Wilmington, MA) born and housed in the Research Support Facility at the University of Kansas Medical Center. Mice were housed on a 12-hour light cycle (600 to 1800 hours) and received water and food *ad libitum*. All research was approved by the University of Kansas Medical Center Institutional Animal Care and Use Committee (protocol number 2016-2344) in compliance with the National Institute of Health Guide for the Care and Use of Laboratory Animals.

### 2.2 Experimental design

Our experimental design is schematized in Figure 1. Briefly, mice underwent NMS or remained unhandled during the first three weeks of life. Voluntary wheel running was introduced at 4-weeks of age, coinciding with a control diet. At 16-weeks of age, half of the sedentary and exercised mice were placed on a HFS diet until 27-weeks of age.

**Figure 1.**
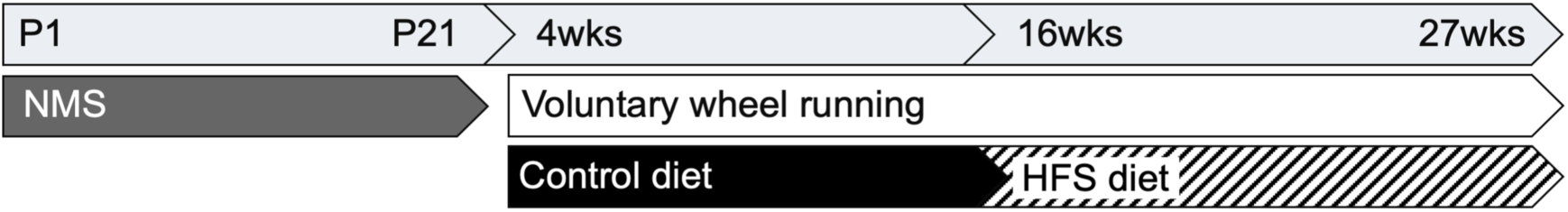
Mice underwent NMS from P1 to P21, or remained unhandled (naïve), and were weaned and pair-housed with littermates on P22. Half of the cages were equipped with a running wheel at 4-weeks of age. All mice were also placed on a control diet at 4-weeks of age. At 16-weeks of age, the control diet was replaced with a HFS diet in half of all cages. Weight gain, distance ran, and diet consumed were measured on a weekly basis. Body composition analysis occurred at 14- and 27-weeks of age. End point measurements including fasting insulin and glucose, glucose tolerance, and adipose analyses were taken at 27-weeks of age.

### 2.3 Neonatal maternal separation

Pregnant C57Bl/6C dams were delivered to the animal facility between 14-16 days gestation. Beginning on postnatal day 1 (P1) until P21, NMS litters were removed *en masse* and placed in a clean glass beaker with bedding from their home cage for 180 minutes (11am-2pm) daily. The beaker was placed in an incubator maintained at 33°C and 50% humidity. Naïve mice remained undisturbed in their home cage except for normal animal husbandry. All mice were weaned on P22 and pair-housed with same sex litter mates. At 4-weeks of age, all mice were placed on a control diet consisting of 20% kcal protein, 70% kcal carbohydrate (3.5% sucrose), 10% kcal fat (3.85 kcal/g; Research Diets, Inc., New Brunswick, NJ; cat. D12110704).

### 2.4 Exercise

At 4-weeks of age, half of the naïve and NMS mice were divided into voluntary wheel running exercised (Ex) or sedentary (Sed) groups. Ex mice were pair-housed with a litter mate in cages equipped with a stainless-steel running wheel (STARR Life Sciences Corp, Oakmont, PA) and Sed mice were pair housed with no access to a running wheel. Distance ran/pair was recorded by STARR Life Sciences VitalView Activity Software version 1.1. Pair housing was implemented to eliminate potential stress associated with single housing of rodents.

### 2.5 High-fat/high-sucrose (HFS) diet

At 16-weeks of age, half of the naïve- and NMS-Sed and -Ex groups were placed on a HFS diet, consisting of 20% kcal protein, 35% kcal carbohydrate (15% sucrose), and 45% kcal fat (4.73 kcal/g; Research Diets), to mimic a Western-style diet.

### 2.6 Food consumption and feed efficiency

Food consumption per pair of mice was measured weekly. Feed efficiency was quantified every week using the following equation: ((weight change per pair)/(calories consumed per pair))*1000. It is presented as the average feed efficiency over the 11-weeks of the experiment after the start of the HFS diet.

### 2.7 Body composition analysis

Mice were weighed and body composition was measured by qMRI using the EchoMRI 1100 (EchoMRI LLC, Houston, TX). Total weight, percent body fat, and free fat mass were quantified every 2-3- weeks. N=4 for all groups except Ex-HFS, n=6.

### 2.8 Glucose Tolerance Test

At 27-weeks of age, following a 6- hour fast, mice were given an intraperitoneal injection of glucose at 1 g/kg body weight. Blood glucose levels were measured via tail clip immediately before glucose injection and 15, 30, 60, and 120-minutes thereafter using a colorimetric assay (PGO enzyme preparation and dianisidine dihydrochloride, Sigma-Aldrich, St. Louis, MO). Area under the curve measurements were calculated using the trapezoidal method [32].

### 2.9 Fasting insulin level

The initial fasting blood collection described above was also used to measure serum insulin levels using an insulin *ELISA* kit according to the manufacturer’s instructions (ALPCO, Salem, NH).

### 2.10 mRNA extraction and RT-PCR

Mice were overdosed with inhaled isoflurane. Epidydimal adipose tissue was dissected, immediately frozen in liquid nitrogen, and stored at −80°C. Frozen tissue was then crushed (Cellcrusher, Portland, OR) and total RNA was isolated using QIAzol Lysis Reagent and the RNeasy Lipid Tissue Mini Kit (Qiagen, Valencia, CA). The concentration and purity were determined using NanoDrop 2000 (Thermo Fisher Scientific, Wilmington, DE) and cDNA was synthesized from total RNA using the iScript cDNA Synthesis Kit (Bio-Rad, Hercules, CA). Quantitative RT-PCR was performed using SsoAdvanced SYBR Green Supermix (Bio-Rad) and a Bio-Rad iCycler IQ real time PCR system with indicated 20uM primers (Table 2; Integrated DNA Technologies, Coralville, IA). Samples were run in triplicate and negative control reactions were run with each amplification series. To reduce variability among efficiency due to fluctuations in baseline fluorescence, the raw PCR data was imported to the LinRegPCR software and PCR efficiency values were derived for each individual sample. Threshold cycle values were subtracted from that of the housekeeping gene PPiB and the fold change over naïve-Sed-Control was calculated using the Pfaffl method [33].

### 2.11 Statistical analysis

Data are reported as mean ± SEM. Calculations of the above measurements were made in Excel (Microsoft, Redmond, WA) and statistical analyses were performed using GraphPad Prism 8 (GraphPad, La Jolla, CA) or IBM SPSS Statistics 24 (IBM Corporation, Armonk, NY). Differences between groups were determined by 2- or 3-way ANOVA or 4-way RM ANOVA and Fisher’s LSD or Bonferroni posttest. Statistical significance was set at *p*<0.05.

## 3. Results

### 3.1 NMS induces greater fat mass

Body weight and composition were assessed at 14-weeks of age, which revealed a significant overall effect of exercise on body weight (Figure 2A, *p*=0.0153), fat mass (Figure 2B, *p*<0.0001), and body fat percentage (Figure 2D, *p*<0.0001), with the latter two measurements having a significant decrease in Ex mice compared to Sed for both naïve and NMS (*p*<0.05). A separate effect of NMS was also observed on fat mass (*p*=0.0223) and percent body fat (*p*=0.0142). No differences were observed in fat-free mass between any groups (Figure 2C, *p*>0.05).

**Figure 2.**
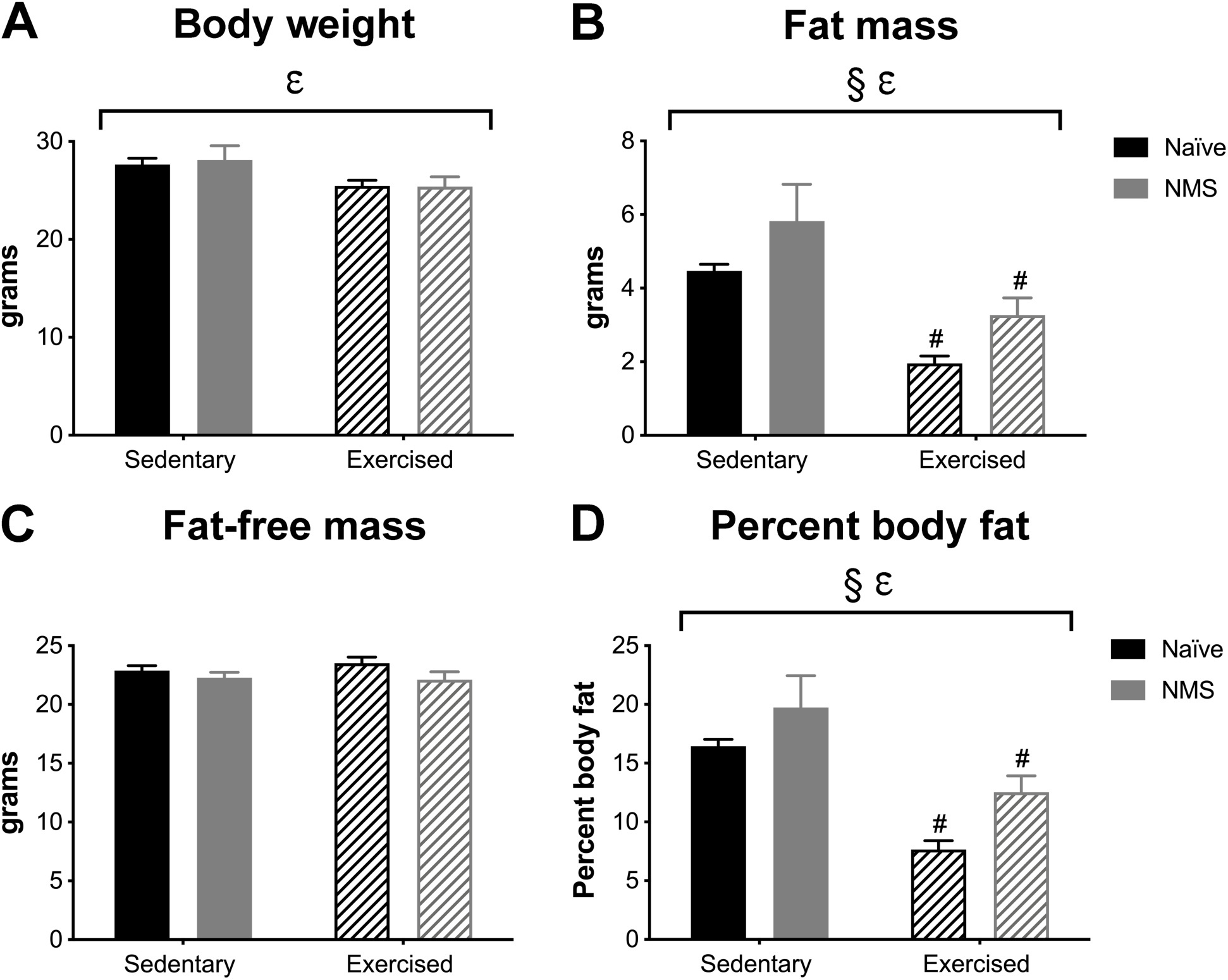
Body weight and composition were measured at 14 weeks of age in naïve and NMS mice that were pair-housed in sedentary caging or given free access to a running wheel beginning at 4-weeks of age. Exercise had a significant impact on body weight (**A**), fat mass (**B**), and percent body fat (**D**). NMS also impacted fat mass (**B**) and percent body fat (**D**). **C**)No group differences were observed in fat-free mass measurements. Brackets indicate a significant effect of exercise NMS (§ *p*<0.05) or exercise (*ε p*<0.05) 2-way ANOVA. # *p*<0.05 Ex vs. Sed, Fisher’s LSD posttest.

### 3.2 Running distance is influenced by stress and diet

Exercised mice were pair housed to avoid additional stress of single housing. Therefore, data are presented as average kilometers ran/day/pair. While on the standard chow diet, NMS mice ran significantly shorter distances per week than naïve mice (Figure 3A, *p*=0.004). Half of the mice were transitioned to a HFS diet at 16 weeks of age. After introduction of a HFS diet, there was a significant impact of NMS (Figure 3A, *p*=0.005) and diet (*p*=0.024) on distance ran per week, with both naïve-HFS and NMS-HFS mice decreasing their running distance by 39.9% and 26.5% at 27-weeks of age, compared to their pre-HFS diet distances, respectively. The total distance ran during the HFS diet intervention was significantly impacted by both NMS (Figure 3B, *p*=0.0013) and diet (*p*=0.0128).

**Figure 3.**
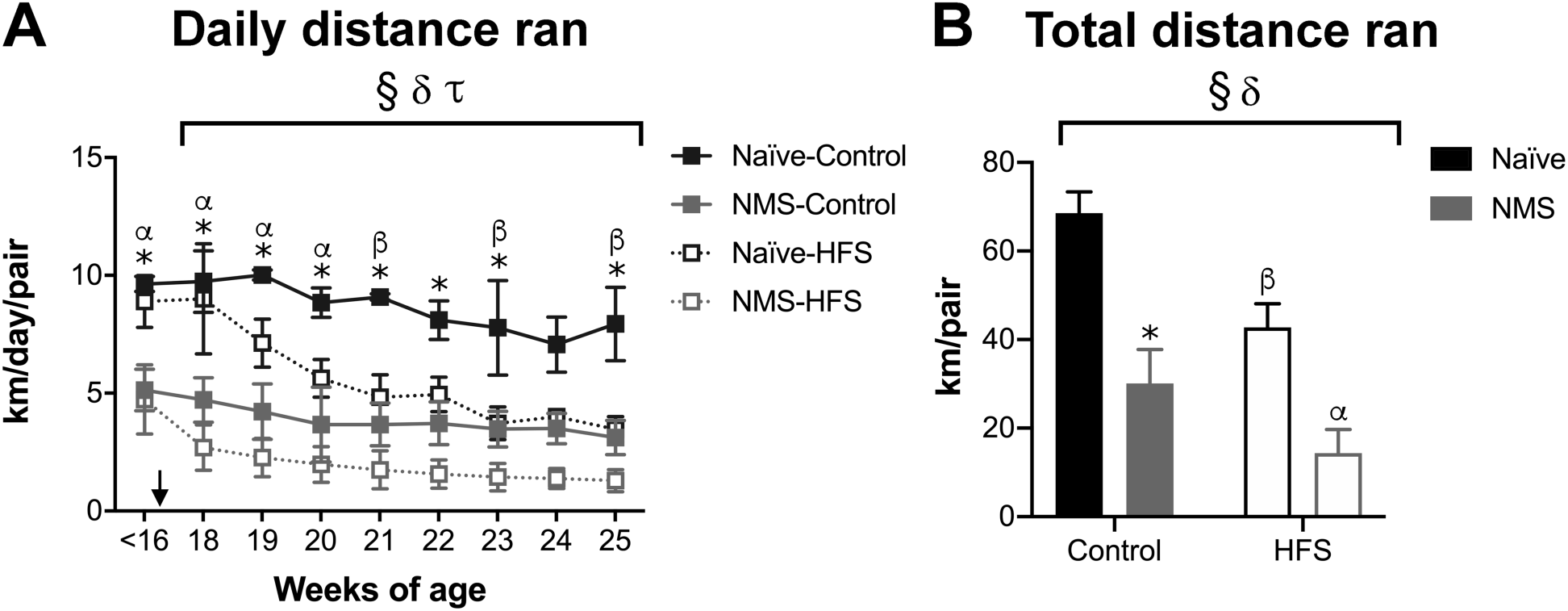
Distance ran/day (km) was calculated for pair-housed naïve and NMS mice. **A**) Prior to 16 weeks of age, at the time of HFS diet introduction (arrow), naïve mice ran significantly farther than NMS mice. Control diet-fed naïve and NMS mice continued to run approximately the same distance/day throughout the experiment, compared to their <16 week distances. Naïve and NMS mice on HFS diet ran significantly shorter distances starting at 20- and 22-weeks of age (*p*<0.05), respectively, compared to their <16-week distances. **B**) Total distance ran for the duration of the diet intervention was significantly impacted by NMS and diet, with both factors reducing distance. All data are reported per pair. Brackets indicate a significant effect of NMS (§ *p*<0.01), diet (*δ p*<0.05), time (*τ p*<0.0001) 3-way RM ANOVA (**A**) or 2-way ANOVA (**B**). * *p*<0.05 naïve-control vs. NMS-control, α *p*<0.05 naïve-HFS vs. NMS-HFS, β *p*<0.05 naïve-control vs. naïve-HFS, Tukey’s multiple comparisons test (**A**) and Fisher’s LSD (**B**).

### 3.3 NMS increases susceptibility to HFS diet-induced weight and fat mass gains

At 27-weeks of age, following 11-weeks on HFS diet, there was a significant overall effect of exercise to reduce body weight (Figure 4A, *p*=0.013), fat mass (Figure 4B, *p*<0.0001) and body fat percentage (Figure 4D, *p*<0.0001). Diet also significantly increased body weight, fat mass, and percent body fat (all *p*<0.0001) with a trend toward impacting fat-free mass (Figure 4C, *p*=0.052). Interaction effects were observed for NMS and exercise (*p*=0.042) and exercise and diet (*p*=0.034) for body fat percentage. In general, HFS groups had significantly greater body weight and fat mass, and had a higher body fat percentage than control diet groups. Exercise attenuated gains in fat mass and percent body fat in naïve- and NMS-control diet mice, however exercise only significantly lowered fat mass (*p*=0.008) and percent body fat (*p*=0.034) in naïve-HFS mice, compared to sedentary HFS-fed mice. Importantly, exercise had no effect on NMS-HFS mice.

**Figure 4.**
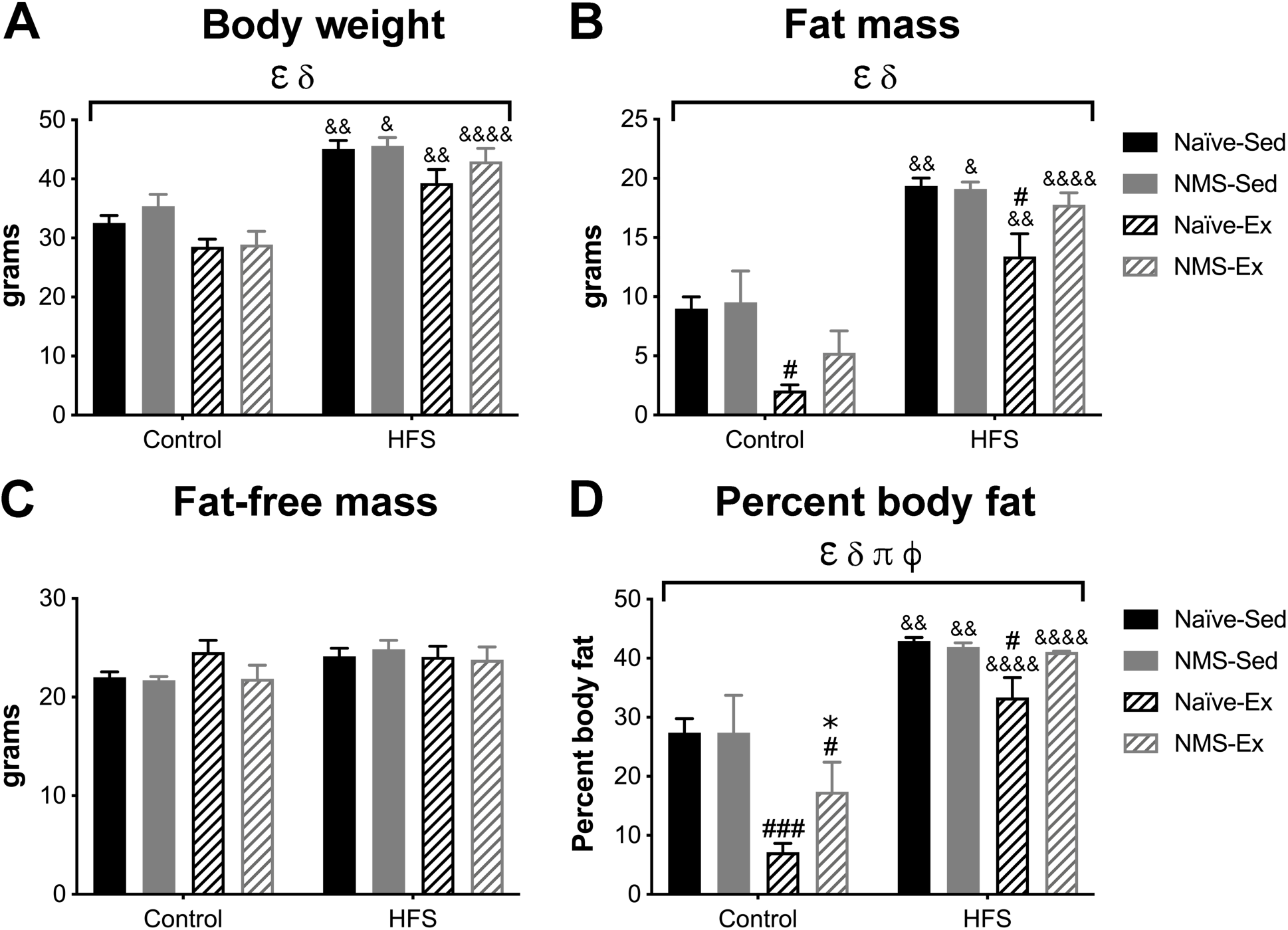
Body weight and composition were measured at 27 weeks of age in naïve and NMS mice that were sedentary or had running wheels and were on a control diet or switched to a HFS diet at 16 weeks of age. Exercise and diet both separately had a significant impact on body weight (**A**), fat mass (**B**), and percent body fat (**D**). A significant interaction effect of NMS and exercise and exercise and diet was also observed on percent body fat. For all three measurements, exercise had a greater impact on HFS diet-fed naïve mice than NMS mice. **C**) Diet showed a trend toward significantly impacting fat-free mass measurements (*p*=0.052). Brackets indicate a significant effect of exercise (*ε p*<0.05), diet (*δ p*<0.0001), NMS/diet interaction (*π p*<0.05), exercise/diet interaction (*ϕ p*<0.05), 3-way ANOVA. * *p*<0.05 vs. NMS vs. naïve, #, ### *p*<0.05, 0.001 Ex vs. Sed, &, &&, &&&& *p*<0.05, 0.01, 0.0001 HFS vs. control, Fisher’s LSD posttest.

### 3.4 Stress, exercise, and diet affect food consumption and feed efficiency

Weekly food consumption, in grams, during the diet intervention, was significantly impacted by NMS (Figure 5A, *p*=0.025), exercise (*p*=0.001), and an NMS/exercise/diet intervention (0.046). Weekly food consumption, in kcal, was also impacted by NMS (Figure 5B, *p*=0.039), exercise (*p*=0.002), and diet (*p*=0.007). For both measures, naïve-Ex mice on control diet ate significantly more than naïve-Sed and NMS-Ex on control diet and naïve-Ex on HFS diet (*p*<0.01). Feed efficiency was significantly impacted by exercise (Figure 5C, *p*=0.012) and diet (*p*<0.0001), with all HFS diet groups having significantly higher feed efficiency than their control-fed counterparts (*p*<0.05), which was significantly reduced in naïve-Ex mice, compared to sedentary (*p*<0.05).

**Figure 5.**
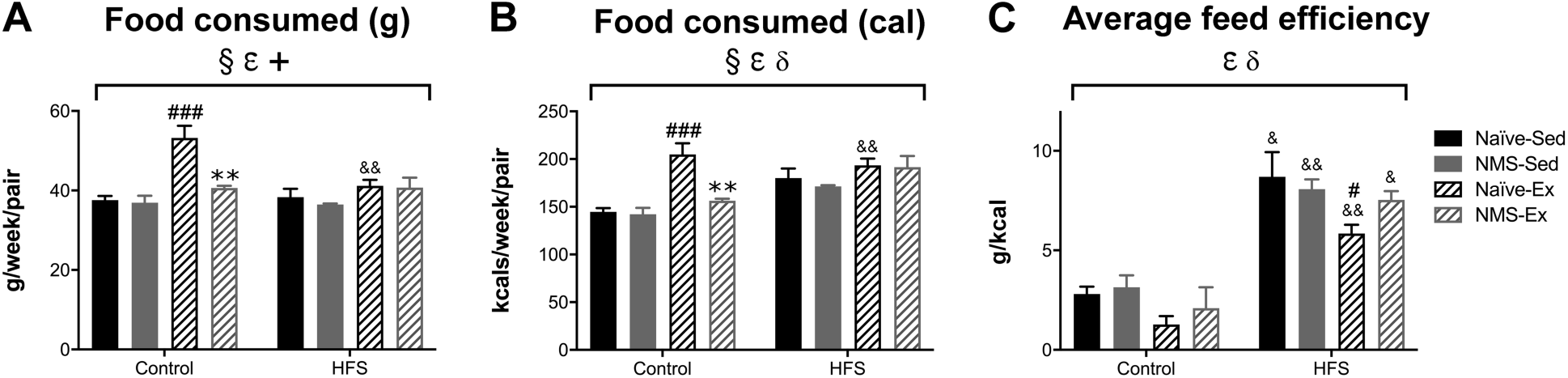
Food consumption per pair was calculated throughout the diet intervention. A significant impact of NMS, exercise, and an NMS/exercise/diet interaction was observed on the amount of food, measured in grams (**A**), was consumed per week. A similar significant impact of NMS and exercise, with an additional impact of diet, was observed on kcal/week consumed (**B**). Naïve-Ex mice on a control diet had the highest consumption, for both measures, compared to control diet-fed naïve-Sed and NMS-Ex mice, and naïve-Ex mice on HFS diet. **C**) Feed efficiency was significantly impacted by exercise and diet, with all HFS diet groups having significantly greater feed efficiency compared to their control counterparts. Exercise significantly attenuated feed efficiency in naïve-HFS mice, whereas it had no impact on NMS feed efficiency regardless of diet. Brackets indicate a significant effect of NMS (§ *p*<0.05), exercise (*ε p*<0.05), diet (*δ p*<0.05), and NMS/exercise/diet interaction (+, *p*<0.05), 3-way ANOVA. ** *p*<0.01 NMS vs. naïve, #, ### *p*<0.05, 0.001 Ex vs. Sed, &, && *p*<0.05, 0.01 HFS vs. control diet, Fisher’s LSD test.

### 3.5 Exercise and diet have opposing effects on fasting insulin level

At 27-weeks of age, mice were fasted for 6-hours and serum insulin level was measured in blood collected by tail-clip. Fasting insulin levels were significantly impacted by exercise (Figure 6A, *p*=0.014) and diet (*p*<0.0001), with sedentary mice on HFS diet having significantly higher serum insulin (*p*<0.01) than control-fed mice, which was significantly reduced by exercise (*p*<0.05).

**Figure 6.**
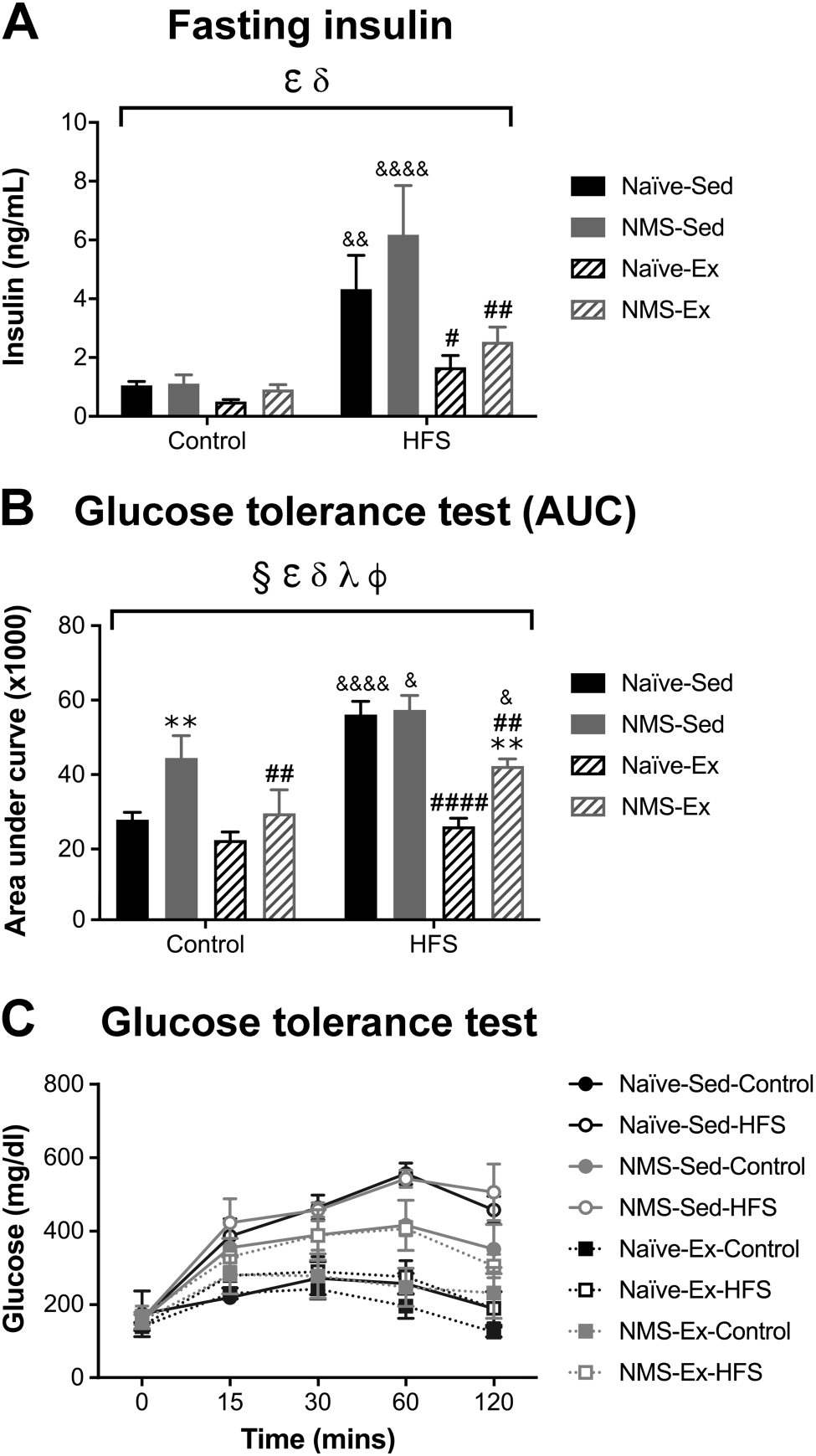
Fasting serum insulin and glucose tolerance were measured at 27-weeks of age. **A**) A significant impact of exercise and diet was observed on fasting insulin levels. HFS diet significantly increased insulin in both sedentary groups, but not in the exercised groups. **B**) A significant effect of NMS, exercise, diet, an NMS/diet interaction, and a diet/exercise interaction was observed on the area under the curve (AUC) measure of blood glucose during the glucose tolerance test. Control-fed NMS-Sed mice had significantly higher blood glucose compared to control-fed naïve-Sed and NMS-Ex mice. All HFS diet-fed groups had significantly increased blood glucose with the exception of naïve-Ex, which was not different from control-fed naïve-Ex mice. Exercise in NMS-HFS mice was moderately protective, as their levels were significantly lower than NMS-Sed-HFS mice, but significantly higher than naïve-Ex-HFS mice. **C**) Individual group measurements at each time point of the glucose tolerance test. Brackets indicate a significant effect of NMS (§ *p*<0.05), exercise (*ε p*<0.05), diet (*δ p*<0.0001), NMS/diet interaction (*λ*, *p*<0.05) and diet/exercise interaction (*ϕ p*<0.05), 3-way ANOVA. ** *p*<0.01 NMS vs. naïve; #, ##, #### *p*<0.05, 0.01, 0.0001 Ex vs. Sed; &, &&, &&&& *p*<0.05, 0.01, 0.0001 HFS vs. control diet, Fisher’s LSD test.

### 3.6 NMS, exercise, and diet influence glucose tolerance

Following an initial blood collection, 1g/kg glucose was administered intraperitoneally and serum blood glucose was measured at 15-, 30-, 60-, and 120-minutes. Both the time course of serum glucose level (Figure 6C) and the area under the curve (AUC; Figure 6B) were analyzed. A significant overall effect of NMS (*p*<0.0001), exercise (*p*<0.0001), diet (*p*<0.0001), an exercise/diet interaction (*p*=0.023) and a NMS/diet/exercise intervention (*p*=0.026) was observed on AUC calculations of serum glucose levels. Interestingly, NMS-Sed mice on control diet had significantly elevated blood glucose, compared to naïve-Sed mice (*p*<0.01), which was significantly lowered in NMS-Ex mice (*p*<0.01). All HFS diet-fed groups had significantly higher blood glucose, compared to control diet groups (*p*<0.05), with the exception of naïve-Ex, which was not comparably different from their control fed counterparts. Exercise was only partially protective of NMS-HFS mice, which had a blood glucose level significantly higher than NMS-control-Ex and naïve-HFS-Ex, but significantly lower than NMS-control-Sed (*p*<0.05). Individual group blood glucose measurements through the test are show in Figure 6C, which further illustrate the lack of impact of exercise in the NMS-HFS diet group.

### 3.7 NMS, exercise, and diet alter epididymal adipose macrophage and mast cell markers as well as neuroendocrine signaling molecules

Epididymal adipose tissue was analyzed for measures of inflammation and neuroendocrine signaling. The mRNA levels for the macrophage markers F4/80 and CD68 were significantly impacted by exercise (Figure 7A, *p*=0.008 and *p*=0.002), diet (*p*<0.0001 for both), and an exercise/diet interaction (*p*=0.003 and *p*=0.006). There was also a significant effect of NMS (*p*=0.005) and a NMS/diet interaction (*p*=0.018) on CD68 mRNA levels. All HFS diet groups had significantly higher F4/80 and CD68 mRNA levels compared to control diet groups (*p*<0.01) with the exception of naïve-Ex-HFS, which was not significantly different from naïve-Ex-Control (*p*>0.05) and was significantly lower than naïve-Sed-HFS (*p*<0.05). Exercise significantly lowered F4/80 and CD68 mRNA levels in HFS-fed NMS mice (*p*<0.05); however, these levels remained significantly higher than those in exercised naïve-HFS fed mice (*p*<0.05). The mRNA levels of CD11b and CD11c were measured to determine macrophage inflammatory status. Diet had a significant impact on CD11b (*p*<0.0001) and CD11c (*p*<0.0001) mRNA levels, and an additional effect of exercise (*p*=0.011) and an exercise/diet interaction (*p*=0.046) was noted for CD11c mRNA levels. Sedentary HFS-fed mice had significantly higher CD11b and CD11c mRNA levels compared to control-fed counterparts (*p*<0.01), which was prevented by exercise for CD11b and partially reversed by exercise for CD11c. It should be noted that the HFS-induced increase in CD11c mRNA levels was 5-15-fold relative to naïve-Sed-control, compared to an increase of 2-3-fold for CD11b.

**Figure 7.**
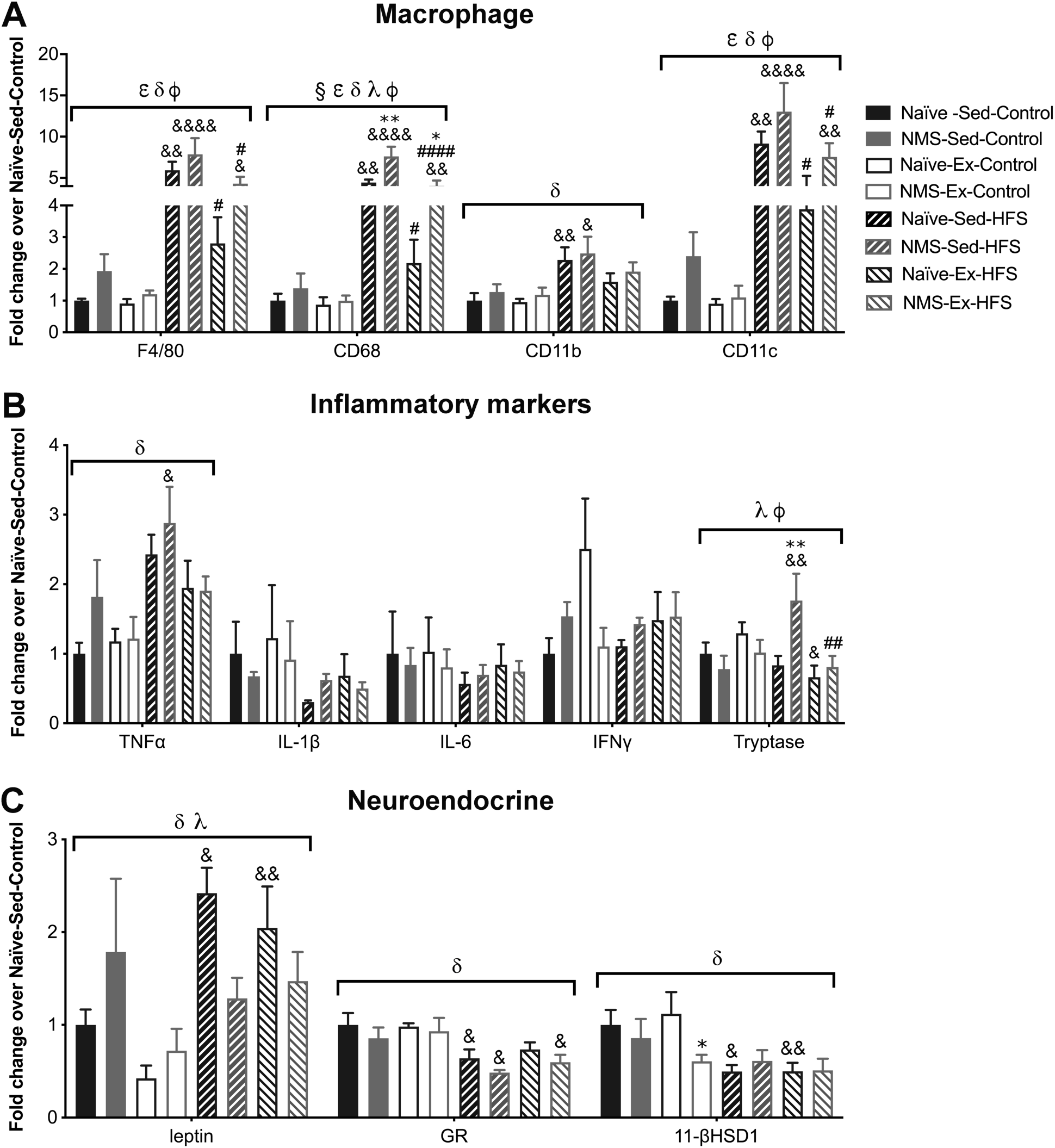
Epididymal adipose tissue was processed for macrophage and inflammatory markers using RT-PCR. **A**) Diet significantly impacted all macrophage genes tested, with additional impacts of NMS on CD68, exercise on F4/80, CD68, and CD11c, NMS/exercise interaction effect on CD68, and an exercise/diet interaction effect on F4/80, CD68, and CD11c. In general, HFS diet increased mRNA levels of all genes in all groups, with the exception of naïve-Ex. The mRNA levels in HFS-fed NMS-Ex mice were significantly lower than NMS-Sed and higher than naïve-Ex. **B**) A significant overall effect of diet was observed on TNFα mRNA levels, with NMS-Sed-HFS having significantly higher TNFα mRNA levels compared to NMS-Sed-Control. Tryptase mRNA levels were significantly impacted by a NMS/diet and exercise/diet interaction, with NMS-Sed-HFS having significantly higher tryptase mRNA levels compared to NMS-Sed-Control, naïve-Sed-HFS, and NMS-Ex-HFS. Naïve-Ex-HFS had significantly lower tryptase mRNA levels compared to naïve-Ex-Control. **C**) A significant overall effect of diet was observed on leptin, GR, and 11-βHSD1 mRNA levels. A significant NMS/diet interaction was also observed on leptin mRNA levels, such that HFS diet only increased leptin mRNA levels in naïve mice, regardless of exercise status, and had no impact on NMS mice. HFS diet also decreased GR and 11-βHSD1 mRNA levels in most groups, compared to their control diet-fed counterparts. Brackets indicate a significant effect of NMS (§ *p*<0.01), exercise (*ε p*<0.05), diet (*δ p*<0.01), a NMS/diet interaction (*λ p*<0.05), or an exercise/diet interaction (*ϕ p*<0.05) 3-way ANOVA; *, ** *p*<0.05, 0.01 NMS vs. naïve, #, ##, #### *p*<0.05, 0.01, 0.0001 Ex vs. sedentary, &, &&, &&&& *p*<0.05, 0.01, 0.0001 HFS vs. control diet, Fisher’s LSD posttest.

The pro-inflammatory cytokine tumor necrosis factor alpha (TNFα) was significantly impacted by diet (Figure 7B, *p*=0.001), particularly in NMS-Sed-HFS mice (*p*<0.05). Interleukin (IL) 1 beta (IL1β), IL6, and interferon gamma (IFN*γ*) mRNA levels were also measured and there were no significant differences found between groups. As a means to evaluate potential mast cell infiltration, the level of tryptase mRNA in epididymal adipose was measured. Significant interaction effects of NMS and exercise (*p*=0.011) and diet and exercise (*p*=0.008) were observed on tryptase mRNA levels, in particular sedentary NMS-HFS fed mice had significantly higher levels compared to both sedentary NMS-control fed (*p*<0.01) and exercised NMS-HFS fed (*p*<0.01) mice. Interestingly, naïve-Ex-HFS had a significantly lower tryptase mRNA level than naïve-Ex-Control mice (*p*<0.05).

To determine potential changes in neuroendocrine signaling, the levels of leptin, glucocorticoid receptor (GR), and 11β-hydroxysteroid dehydrogenase type 1 (11βHSD1) mRNA were measured (Figure 7C). Diet (*p*=0.007) and a NMS/diet interaction (*p*=0.019) significantly impacted leptin mRNA levels. The impact of diet was largely observed only in naïve mice, as these groups had significantly increased leptin mRNA compared to their control diet counterparts, regardless of exercise (*p*<0.05). In contrast, HFS diet had no significant impact on leptin mRNA levels in NMS mice (*p*>0.05). Diet also significantly impacted GR (*p*<0.0001) and 11βHSD1 mRNA levels (*p*=0.001). All HFS groups, with the exception of naïve-Ex, had a significantly lower GR mRNA level compared to control diet mice (*p*<0.05) and naïve-Sed-HFS and -Ex-HFS mice had a significantly lower 11βHSD1 mRNA level compared to control diet mice (*p*<0.05). Finally, NMS-Ex-Control mice had a significantly lower 11βHSD1 mRNA level compared to naïve-Ex-Control (*p*<0.05).

## 4. Discussion

Obesity-related MetS is highly prevalent in today’s society and dramatically increases the risk for the development of type 2 diabetes and cardiovascular disease. In humans, environmental factors including early life stress [3–5], poor diet [16, 17], and low activity level [34–36] increase one’s risk to develop the MetS. Many pre-clinical studies have investigated the influence that HFS diet [37–40] and exercise [31, 41, 42] have on the development of obesity-related MetS, while only a few have investigated the impact of early life stress [14, 15]. To our knowledge, this is the first study to combine early life stress, exercise, and diet to evaluate outcomes related to this disorder.

The primary findings showed that 14-week old male mice that had undergone NMS had similar body weight, but increased body fat percentage, compared to naïve mice, regardless of caging conditions. *Ad libitum* access to a HFS diet predictably increased body weight and body fat percentage in both naïve and NMS sedentary mice. However, daily exercise, via voluntary wheel running, only positively impacted body fat percentage in HFS diet-fed naïve mice, but not NMS mice who had undergone early life stress. These findings are similar to other studies investigating the effect of early life stress exposure on diet-induced metabolic outcomes. For example, Yam et al., [15] used the limited nesting material paradigm of early life stress and demonstrated that although stressed mice exhibited comparably reduced weight and adipose tissue at weaning, the stressed mice were more susceptible to high-fat diet-induced obesity than non-stressed mice. Additionally, Murphy et al., [43] used maternal separation with early weaning as a stress paradigm and demonstrated that stressed adult male and female mice on a low-fat diet had increased fat mass compared to non-stressed mice. Interestingly, female stressed mice were more susceptible to high-fat diet-induced changes in body weight and fat compared to non-stressed mice, while male stressed mice consuming a high-fat diet had similar body weight and fat compared to non-stressed mice. We are currently investigating potential sex differences in diet-induced metabolic outcomes in our NMS model.

Our results suggest that exercise was more beneficial in preventing weight and body fat gain in naïve mice compared to NMS mice. Both reduced running volume and a more disrupted metabolic phenotype induced by NMS may underlie these findings. Interestingly, naïve-Ex mice ate significantly more control diet compared to the other groups, yet they demonstrated appropriate energy balance by running more than NMS-Ex mice. The reduced running distance, and putatively reduced energy expenditure by NMS mice, could partially underlie their increased weight and body fat gain compared to naïve mice. However, differences in feed efficiency suggest that basal differences in metabolism may also be at play. Exercise significantly lowered feed efficiency in HFS diet-fed naïve groups when compared to sedentary counterparts. In contrast, exercise had no impact on feed efficiency in NMS mice on either a control or HFS diet. Again, the low running distance of the NMS, particularly those on HFS diet, likely contributes to the moderate impact on feed efficiency. We used voluntary wheel running because this form of exercise is considered rewarding to rodents as opposed to forced treadmill running, which is considered a stressor and can cause anxiety like behaviors [44–46]. We have previously observed an attenuation of bladder hypersensitivity and mast cell degranulation in female mice of the same NMS model [47]. The NMS mice in this study also ran approximately half the weekly distance of naïve mice, suggesting this is a consistent phenotype following NMS. Future studies will incorporate limiting daily running wheel distances or the use of treadmills to determine the impact of NMS in mice with equivalent aerobic output.

Impaired glucose homeostasis is one symptom of MetS and a hallmark of prediabetes [25]. The sedentary, control diet-fed NMS mice in this study exhibited glucose intolerance, which was significantly improved by exercise. This finding is in line with other groups that have demonstrated the positive effect that exercise has on glucose tolerance in rodents [41, 48, 49]. Exercise prevented glucose intolerance in naïve mice on HFS diet, whereas exercised NMS mice on the HFS diet only had moderately, yet significantly, improved glucose tolerance, which was still higher than that of similarly-treated naïve mice. The only HFS-fed group that did not develop glucose intolerance was the naïve-Ex group. Another intriguing finding was that NMS induced glucose intolerance on a control diet, supporting the epidemiological association between early life stress in humans and the development of obesity and related metabolic disorders in adulthood [2–5]. In direct relation to the glucose homeostasis data, fasting insulin was also impacted by NMS, exercise, and diet. Accordingly, the sedentary NMS mice on a HFS diet in our study had the highest fasting insulin level.

White adipose tissue distribution and composition is important in predicting the development of obesity-related diseases because it plays a key role in regulating systemic metabolic function and inflammation [50]. Visceral fat accumulation is associated with a greater risk of metabolic dysfunction [51] and leads to chronic low-grade inflammation [52]. Low-grade inflammation is, in turn, associated with increased pro-inflammatory macrophage and MC infiltration as well as the increased prevalence of crown-like structures and has been demonstrated in both obese mice and humans [53–56]. Our results indicate that HFS diet and NMS significantly increased pro-inflammatory macrophage markers in epididymal adipose tissue. While exercise did significantly reduce these levels overall, this was predominantly observed in the naïve-HFS mice, whereas the levels in NMS-Ex-HFS mice were still significantly elevated. Similarly, TNFα mRNA levels were also significantly increased by HFS diet. This is a mechanistically significant finding because TNFα affects insulin sensitivity [57]. Interestingly, the group in our study with the highest TNFα in epididymal adipose, NMS-Sed-HFS, also had the highest fasting insulin level, suggesting that insulin resistance could be at least partially driven by increased visceral adipose TNFα.

Gurung et al., [58] recently showed that subcutaneous adipose tissue of individuals with MetS had a 2.5-fold increase in MCs compared to control subjects. This increase in MCs was positively correlated with markers of fibrosis and inflammation. Obese subjects have also been shown to have significantly higher serum tryptase levels [59]. Furthermore, high-fat diet-induced obese mice display increased MCs in visceral adipose tissue [56]. Here, we observed significant NMS/diet and exercise/diet interactions on tryptase mRNA levels in epididymal adipose. By far, the highest level of tryptase mRNA was found in NMS sedentary HFS adipose, which was reduced by exercise. These results imply that, much like with glucose intolerance and insulin levels, the combination of early life stress, sedentary caging, and a HFS diet had a cumulative effect on increasing epididymal adipose MC infiltration.

White adipose tissue is a complex endocrine organ that secretes adipokines, including leptin [60], which communicates with its receptor in the hypothalamus to regulate food intake and energy expenditure by shutting down hunger signals [61]. This circuitry is not fully programmed at birth making it susceptible to the influence of environmental factors [62]. Circulating leptin usually peaks during development from P4-P16 in rodents [63] and this peak is important for proper formation of the hypothalamic feeding circuitry [61]. Early life stress has been shown to reduce leptin levels early in life [64, 65], which could lead to altered programming of the leptin-hypothalamic circuit and long-term changes in hunger signals and energy expenditure. Our results suggest that NMS alters leptin signaling. Naïve mice on the HFS diet had significantly increased leptin levels compared to mice on the control diet, which is expected so that hunger signals are decreased and energy expenditure is increased to compensate for the extra energy being consumed. However, NMS mice on the HFS diet did not show this increase. Additionally, NMS sedentary mice on the control diet displayed increased leptin mRNA levels in adipose, which is not expected on a control diet and suggests that these mice may be leptin resistant. It has been hypothesized that a hyperactive HPA axis, which leads to the elevation of GC concentrations, is involved in leptin resistance in obesity [66]. Our NMS protocol involves separating mice from P1-P21, which includes the important window of development of the leptin-hypothalamic feeding circuitry [61, 63]. Therefore, proper programming of both the HPA axis and leptin signaling could be affected by NMS and have long-term consequences on metabolism.

Increased circulating glucocorticoid levels have been associated with the development of obesity, particularly in the visceral region [67]. This is likely due to glucocorticoid-induced differentiation of pre-adipocytes into mature adipocytes and increased lipoprotein lipase activity, which can increase the amount of adipose tissue and subsequently cause weight gain [62]. Clinical studies in obese patients have shown simultaneous increases in GR and 11βHSD1 [68, 69], which converts inactive glucocorticoids to their active form [70]. However, pre-clinical research showed that a high-fat diet actually caused a decrease in adipose 11βHSD1 [71, 72]. Similarly, we found that there was an overall reduction in 11βHSD1 and GR mRNA in epididymal adipose tissue due to HFS diet. We hypothesize that this down-regulation of 11βHSD1 and GR could be an adaptive mechanism to circumvent the detrimental metabolic consequences of increased GC signaling. This adaptive mechanism appears to function better in our naïve mice on a HFS diet as their 11βHSD1 mRNA levels were significantly reduced while NMS mice 11βHSD1 mRNA levels did not reach significance.\

Overall, exercise attenuated obesity-related MetS outcomes in both naïve and NMS mice, but appeared to have a stronger effect in naïve mice, most likely due to differential running distances between groups. We have previously reported on reduced running distances by female NMS mice [47], and have consistently observed this behavior among multiple cohorts of mice (unpublished observation). Although running distance was significantly lower, we have observed an attenuation of urogenital hypersensitivity and bladder overactivity due to voluntary wheel running in female NMS mice [47]. We hypothesize that differences in running distance could be due to changes in reward pathways, as we have also observed gene expression changes indicative of reduced hippocampal regulation of the HPA axis [27, 28, 47, 73]. Lovallo [74], explains that early life stress in humans can lead to altered dopaminergic signaling, which influences motivated behaviors [75], including physical activity. Our future studies will be aimed at determining the underlying mechanism driving reduced voluntary wheel running in NMS mice.

In summary, NMS, exercise, and diet all influence factors related to MetS. This was first demonstrated by our finding that chow-fed NMS sedentary male mice have altered body composition compared to naïve and Ex mice and was further exemplified by our long-term HFS diet study. The NMS-Sed-HFS mice had the highest fasting insulin level, the highest mRNA levels of TNFα, the pro-inflammatory macrophage marker CD11c, and MC tryptase in epididymal adipose tissue, suggesting this group was more prone to visceral inflammation and development of insulin resistance compared to the other groups. Our results also implied that our NMS-Sed mice on control chow were also more susceptible to developing symptoms of MetS compared to naïve- or NMS-Ex-control mice. They had increased TNFα and CD11c mRNA levels, compared to other groups on the control diet, as well as increased GTT AUC, which implies they are more prone to visceral inflammation and are less glucose tolerant. Taken together, these data suggest that early life stress exposure reduces voluntary running distance and the efficiency of exercise to attenuate the negative effects of a HFS diet.

## Acknowledgments

The authors would like to thank Drs. Isabella Fuentes, Paige Geiger, Kenneth McCarson, Andrea Nicol, and Doug Wright for their thoughtful contributions towards the development, execution, and interpretation of this project. We would also like to acknowledge Ruipeng Wang and Dr. Xiaofang Yang for their technical assistance in carrying out the described experiments.

